# DNA polymerase η is regulated by mutually exclusive mono-ubiquitination and mono-NEDDylation

**DOI:** 10.1101/2024.10.12.618026

**Authors:** Natália Cestari Moreno, Emilie J. Korchak, Marcela Teatin Latancia, Dana A. D’Orlando, Temidayo Adegbenro, Irina Bezsonova, Roger Woodgate, Nicholas W. Ashton

## Abstract

DNA polymerase eta (Pol η) is a Y-family translesion polymerase responsible for synthesizing new DNA across UV-damaged templates. It is recruited to replication forks following mono-ubiquitination of the PCNA DNA clamp. This interaction is mediated by PCNA-interacting protein (PIP) motifs within Pol η, as well as by its C-terminal ubiquitin-binding zinc finger (UBZ) domain. Previous work has suggested that Pol η itself is mono-ubiquitinated at four C-terminal lysine residues, which is dependent on prior ubiquitin-binding by its UBZ domain. Here, we show that Pol η can be modified at the same lysine residues by the ubiquitin-like protein, NEDD8. Like ubiquitination, this modification is driven by non-covalent interactions between NEDD8 and the UBZ domain. While only a small proportion of Pol η is mono-NEDDylated under normal conditions, these levels rapidly increase by inhibiting the COP9 signalosome, suggesting that mono-NEDDylation is maintained under strong negative regulation. Finally, we provide data to support that mono-ubiquitination is important for Pol η foci formation and suggest that NEDDylation disrupts this process. These results reveal a new mechanism of Pol η regulation by ubiquitin-like proteins.

## INTRODUCTION

DNA replication is an essential process that must be completed each cell cycle prior to chromosome segregation. The bulk of DNA synthesis is catalyzed by replicative polymerases of the B-family - DNA polymerases alpha (Pol α), epsilon (Pol ε) and delta (Pol δ) (1). These enzymes replicate DNA with near-perfect accuracy, in part due to their closed active sites that accommodate only correct Watson-Crick base pairs (2). This accuracy however comes at an expense, as these polymerases are largely unable to synthesize new DNA from damaged templates. Replicative DNA synthesis is therefore vulnerable to disruption by DNA damage adducts. One mechanism that cells use to overcome replication barriers is to employ translesion synthesis (TLS) by Y-family DNA polymerases. These specialized polymerases have comparatively open active sites, allowing them to bypass DNA lesions, such as UV-induced photoproducts (3). Y-family DNA polymerases exhibit a remarkable degree of conservation across all domains of life, underscoring their fundamental roles in preserving genome integrity (4). While the catalytic domains of each polymerase is structurally similar, each is thought to be tailored to bypass specific damage types, often referred to as their “cognate lesions” (5). For example, Pol η contains a notably expansive active site, allowing it to accommodate pyrimidine dimers and corresponding purine deoxynucleotides (6,7). The important role of Pol η in bypassing DNA photodamage is evident in the cancer-prone syndrome, Xeroderma Pigmentosum Variant (XP-V), which results from deleterious Pol η mutations. Cells from XP-V patients effectively remove DNA damage through excision repair mechanisms, although exhibit deficient DNA synthesis following exposure to ultraviolet (UV) irradiation (6). As a result, XP-V patients are at a heightened risk of developing skin cancer in body areas exposed to sunlight.

While TLS polymerases are important enzymes for tolerating DNA damage, their permissive active sites – which are necessary to accommodate DNA lesions – compromise their accuracy in inserting new nucleotides. Employing TLS therefore carries the risk of unintended mutagenesis (4). Because of this, TLS polymerases are tightly regulated to prevent their inappropriate use (8). One way this is achieved is through the post-translational modification of pathway proteins. This is exemplified by mono-ubiquitination of the PCNA sliding clamp in response to DNA damage (9). While PCNA is an important interactor of all replicative polymerases, its mono-ubiquitination generates a binding platform for TLS polymerases to engage with the replication fork. This is due to the presence of one or more ubiquitin-binding domains in the C-termini of each Y-family DNA polymerase, which can bind directly to the modified form of PCNA (10–12). Aside from ubiquitin, many ubiquitin-like (UBL) proteins also have key regulatory roles in the cell, including SUMO-1/-2/-3, ISG15 and NEDD8 (13). Although the primary sequences of these proteins vary, all contain a near-identical ubiquitin-like fold and can be conjugated to lysine residues of substrate proteins to regulate a wide range of cellular functions. NEDDylation has been increasingly recognized as an important form of regulation for the DNA repair pathways. Indeed, NEDD8 molecules have been found to localize to sites of DNA damage in cells treated with a range of DNA-damaging agents (14). Relatively few specific substrates of NEDDylation in these pathways have however been identified.

Previous work has found that Pol η is modified by both mono- and poly-ubiquitination. While poly-ubiquitination regulates Pol η protein turnover through the ubiquitin-proteasome system (15), mono-ubiquitination at one of four lysine residues on the Pol η C-terminus is thought to regulate its interaction with other proteins (10,16,17). In this work, we demonstrate that Pol η is also a substrate of NEDD8. Like mono-ubiquitination, mono-NEDDylation is regulated by the ability of Pol η to noncovalently bind NEDD8 via its UBZ domains, directing NEDD8 conjugation to the Pol η C-terminus. This modification is mutually exclusive with ubiquitination and negatively regulates Pol η accumulation at sites of DNA damage. These results thereby demonstrate a new mechanism of Pol η regulation by NEDD8.

## MATERIAL AND METHODS

### Expression vectors

Mammalian and bacterial expression vectors created for this work are available from Addgene and are summarized in Table S1. Plasmid files are also available from Mendeley Data. Synthetic DNA fragments were chemically synthesized by GenScript (Piscataway, NJ).

#### Mammalian expression vectors

The wild-type HA Pol η expression vector (pJRM56) has been described previously (18) (Addgene # 201671), as has the wild-type FLAG Pol η vector (pJRM160) (19) (Addgene # 221897). Vectors expressing FLAG Pol η point mutations or deletions, or FLAG Pol η ubiquitin/NEDD8 fusion proteins, were created by modifying the wild-type FLAG Pol η vector. This was achieved by synthesizing new gene fragments and subcloning into the wild-type plasmid using restriction sites flanking or contained within the Pol η coding sequence. GFP tagged Pol η was expressed from pEGFP-C1 (Clontech), that we firstly modified by addition of an SV40 nuclear localization signal (ATG<u>CCAAAGAAGAAGCGAAAGGTA</u> GCAGATCCA) upstream of the EGFP coding sequence. WT and mutant Pol η coding sequences were cloned downstream of EGFP using the XhoI - BamHI restriction sites.

The HA ubiquitin plasmid has been described previously (Addgene # 131258) (20). The wild-type HA NEDD8 expression vector was created by cloning a chemically synthesized coding sequence of NEDD8 into the KpnI - BamHI restriction sites of pcDNA3.1(+)-N-HA (Genscipt).

#### Bacterial expression vectors

The pET15b-Pol η UBZ (residues 628-662) expression vector (21) was a kind gift from Pei Zhou (Duke University Medical Center). His-ubiquitin was expressed from a previously described pET-15b (Sigma-Aldrich) plasmid (22). The His-NEDD8 plasmid was created by subcloning a codon optimized NEDD8 coding sequence into the NdeI – BamHI restriction sites of pET-28b(+) (MilliporeSigma).

### Mammalian Cell Culture

293T cells were obtained from ATCC (cell line # CRL-3216). Lab stocks of MRC-5 SV2 cells (23) (abbreviated MRC-5) were authenticated by STR profiling (ATCC # 135-XV). Both cell lines were tested for mycoplasma contamination by PCR (ATCC # 30-1012K). Cell lines were cultured in Dulbecco’s Modified Eagle Medium High Glucose (DMEM, Thermo Fisher Scientific # 11965118) supplemented with 10% fetal bovine serum (FBS, Gibco, Thermo Fisher Scientific # A5670701), and 1% penicillin-steptomycin (10,000 units mL^-1^ penicillin, 100mg mL^-1^ streptomycin; Gibco, Thermo Fisher Scientific # 15140122). Cells were cultured in a controlled environment that was maintained at 370°C with a humidified atmosphere containing 5% CO_2_.

### Inhibitors and DNA damage induction

The NEDD8 activating enzyme (NAE) inhibitor, MLN4924, was purchased from Selleckchem (# 905579-51-3), dissolved in DMSO to a concentration of 10 mM, and stored in aliquots at −80 °C. Working solutions were prepared by diluting stock solutions further with DMSO to 1 mM. 293T cells were treated with 0.1 or 0.3 μM MLN4924 for 16 hours prior to harvesting.

CSN5i-3 was purchased from Biotechne (# 7089/2), dissolved in DMSO to a concentration of 1 mM, and stored in aliquots at −20 °C. 293T cells were treated with 1 μM CSN5i-3 for 2 hours prior to harvesting or further treatment.

DNA damage was induced with a UVC germicidal lamp (254 nm). Prior to UVC exposure, the culture medium was replaced with phosphate-buffered saline (PBS). For both the immunoprecipitation and immunofluorescence assays, cells were irradiated with 20 J m² of UVC. After irradiation, PBS was replaced with culture media, and the cells were collected according to the protocol of each experiment.

### Immunoprecipitation

The immunoprecipitation assays were performed as previously described (20,22). Briefly, 293T cells were resuspended and sonicated in immunoprecipitation buffer (20 mM HEPES pH 7.5, 150 mM KCl, 5% glycerol, 10 mM MgCl_2_, 0.5% Triton X-100) supplemented with 1 x protease inhibitor cocktail, 50 μM PR-619 (Selleckchem #S7130) and Pierce Universal Nuclease for Cell Lysis (1:5000, Thermo Fisher Scientific #88700). Magnetic anti-FLAG M2 beads (Sigma-Aldrich # M8823) were used for the immunoprecipitation of FLAG-tagged proteins. In all cases, conjugated beads were washed in immunoprecipitation buffer, then incubated with whole cell lysates for 1 hour at 4 °C. Beads were then washed 4 x with immunoprecipitation buffer, 2 x with immunoprecipitation buffer modified to contain 250 mM KCl and then proteins eluted by incubating the beads in 100 mM pH 2.3 glycine on a shaker for 10 min, followed by neutralization of the sample with 500 mM Tris pH 7.4.

### Immunoblotting

Immunoblotting was performed using a standard protocol (20). Briefly, samples were separated by electrophoresis on a 15-well 1.5 mm 4–12% Bis-Tris NuPage precast gel (Thermo Fisher Scientific) and transferred to nitrocellulose membranes prior to incubation with the following primary antibodies: Rb-α-NEDD8 (E19E3; CST #2754), Ms-α-FLAG (M2; MilliporeSigma #F1804), Rb-α-HA (MilliporeSigma #3724S), Rb-α-PCNA (Cell Signaling #D5C7P), Ms-α-PCNA (Santa Cruz #K3023), Rb-α-actin (Cell Signaling #5057S), Ms-α-H3 (Cell Signaling #8173S). Primary antibodies were detected using IRDye 680RD or 800CW-conjugated donkey anti-mouse or anti-rabbit fluorescent secondary antibodies (Li-Cor) and visualized using an Odyssey CLX infrared imaging system (Li-Cor). Immunoblots were quantified using Image Studio software (Li-Cor). Relative mono-ubiquitination and mono-NEDDylation of FLAG Pol η were calculated as described in the figure legends.

### Protein expression and purification

The UBZ domain of Pol η (amino acids 628–662), NEDD8 and ubiquitin were expressed and purified in a similar manner. All three plasmids were transformed into *Escherichia coli* BL21(DE3) cells. Unlabelled proteins were expressed in 1 L of Luria broth (LB) media and ^15^N-labelled proteins were expressed in M9 minimal media supplemented with 20 μM Zn^2+^ and containing ^15^NH_4_Cl as a sole source of nitrogen. Transformed cells were grown at 37°C until OD_600_ of 0.8–1.0. Protein expression was induced with 1 mM Isopropyl β-D-1-thiogalactopyranoside (IPTG) overnight at 20°C.

Cells were harvested by centrifugation, resuspended in a buffer containing 20 mM phosphate buffer pH 8, 250 mM NaCl, 10 mM imidazole, 1 mM PMSF, lysed by sonication, and centrifuged at 15,000 rpm for 1 hour. The supernatant was filtered and applied to a TALON HisPur cobalt resin (Thermo Scientific #89964). Proteins were eluted in a buffer containing 20 mM phosphate buffer pH 8, 250 mM NaCl and 300 mM imidazole. Thrombin protease was then added to samples overnight at 4°C to remove the 6-His tag. Proteins were subjected to size-exclusion chromatography using a HiLoad Superdex 75 column (Cytiva) in a buffer containing 25 mM Na_2_HPO_4_ pH 7.0, 100 mM KCl and 2 mM dithiothreitol (DTT).

### NMR titration experiments

All NMR experiments were collected at 800 MHz (^1^H) Bruker Avance Neo spectrometer, equipped with a cryogenic probe, at 25°C. 150 μM ^15^N sample of Pol η UBZ domain was gradually titrated with either unlabelled NEDD8 or ubiquitin to a final molar ratio of 1:3 for UBZ:NEDD8 and 1:5 for UBZ:ubiquitin. 100 μM ^15^N NEDD8 was gradually titrated with unlabelled ubiquitin to a final molar ratio of 1:12 (NEDD8:UBZ). ^1^H-^15^N HSQC spectrum was collected for each of the titration points to follow binding.

Data was processed using NMRPipe (24) and analysed using Sparky (25). All data processing and analysis were performed on the NMRbox (26) platform. Per-residue NMR chemical shift perturbations (Δω_obs_) were calculated using the equation: Δω_obs_ = (Δω_N_^2^ + Δω_H_^2^)^1/2^ where Δω_N_ and Δω_H_ are the chemical shift differences between free and bound samples measured in Hz for ^15^N and ^1^H, respectively. The obtained Δω_obs_ values in ^1^H ppm were mapped onto the structure of UBZ (PDB: 3WUP) (27) and NEDD8 (PDB: 1NDD) (28) to reveal the binding interfaces.

NMR titrations were also used to determine UBZ:NEDD8 and UBZ:ubiquitin binding affinities. Only non-overlapping peaks were selected for analysis. The Δω_obs_ were plotted as a function of ligand concentration and the dissociation constants (K_D_) were determined by fitting the curves in GraphPad Prism 10.3.1 using the following equation:

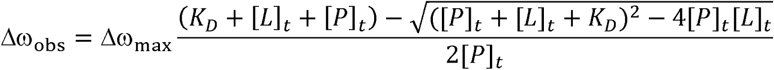

Where [P]_t_ and [L]_t_ are the total protein and ligand concentrations and Δω_max_ is chemical shift difference at saturation.

### Molecular docking

The structural model of the UBZ domain in a complex with NEDD8 was generated with HADDOCK 2.4 (High Ambiguity Driven Docking) using experimental NMR chemical shift perturbations. NEDD8 residues T7, L8, I13, Y45, K48, Q49, H68, L69, V70, and L71 and the Pol η UBZ domain residues W645, D652, F655, and A656 which undergo the largest NMR chemical shift perturbations upon binding were chosen to define the interface. Previously reported structures of NEDD8 (PDB: 1NDD) and the UBZ domain (PDB: 3WUP) were used for docking. Docking yielded 123 structures in 10 clusters. The top cluster with the lowest z-score was chosen as the most reliable.

### Protein structure prediction

AlphaFold3 and the AlphaFold Server (29) were used to model predicted interactions between PCNA and the C-terminus of Pol η, the Pol η C-terminus and ubiquitin, and the Pol η UBZ and ubiquitin. Models were visualized in Pymol V2.5.8 (Schrödinger) using the top-ranking predictions per seed.

### Foci Formation

MRC-5 cells were seeded at equal densities into 60 mm dishes, each containing two 10 mm coverslips, and incubated overnight to allow for attachment. The following day, cells were transfected with 30 µL of TurboFectin, 250 µL of Opti-MEM, and 10 µg of plasmid DNA. After 24 hours, cells were irradiated with 20 Jm² of UV-C and harvested six hours post-irradiation. Coverslips were then recovered, washed with cold PBS, fixed in 4% paraformaldehyde, rinsed with deionized water, and mounted on slides using ProLong Gold antifade reagent containing DAPI. Imaging was conducted using a ZEISS Axiolab 5 fluorescence microscope with a 63x objective lens. Image analysis was performed using Fiji software. The DAPI-stained images were converted to 16-bit grayscale, thresholds were adjusted to generate binary masks, and the Watershed algorithm was applied as necessary. Particle analysis settings were adjusted to outline particles and exclude those on the edges. In the EGFP channel, images were processed to identify foci by converting to 16-bit grayscale, using nuclei masks from the DAPI images to identify transfected cells, and applying the ‘Find Maxima’ function. New images displaying points of maxima were generated and analyzed using the ROI Manager tool to accurately measure and record the foci as Raw Integrated Density (RawIntDen), with the number of foci per nucleus calculated by dividing this value by 255. Statistical analysis was performed using GraphPad Prism 10.2.2. An ordinary one-way ANOVA followed by a post-hoc multiple comparisons test was conducted to compare the groups, with each group consisting of more than 100 transfected cells.

## RESULTS

### Pol η is a substrate of NEDD8

NEDD8 is a known regulator of DNA repair and replication, although few NEDD8 targets in these pathways have been identified. Two recent whole proteome mass-spectrometry based screens have however revealed several hundred endogenous proteins modified by NEDD8, including a small number involved in DNA repair (30,31). Of these, we were interested to note that both screens identified peptides mapping to the C-terminus of Pol η. Furthermore, NEDDylation was detected at two lysine residues – K682 and K709 – that are also known to be sites of mono-ubiquitination (**Figure 1A**) (16,17). We therefore considered that Pol η might also be regulated by NEDD8, potentially via a competitive mechanism with mono-ubiquitination.

**Figure 1:**
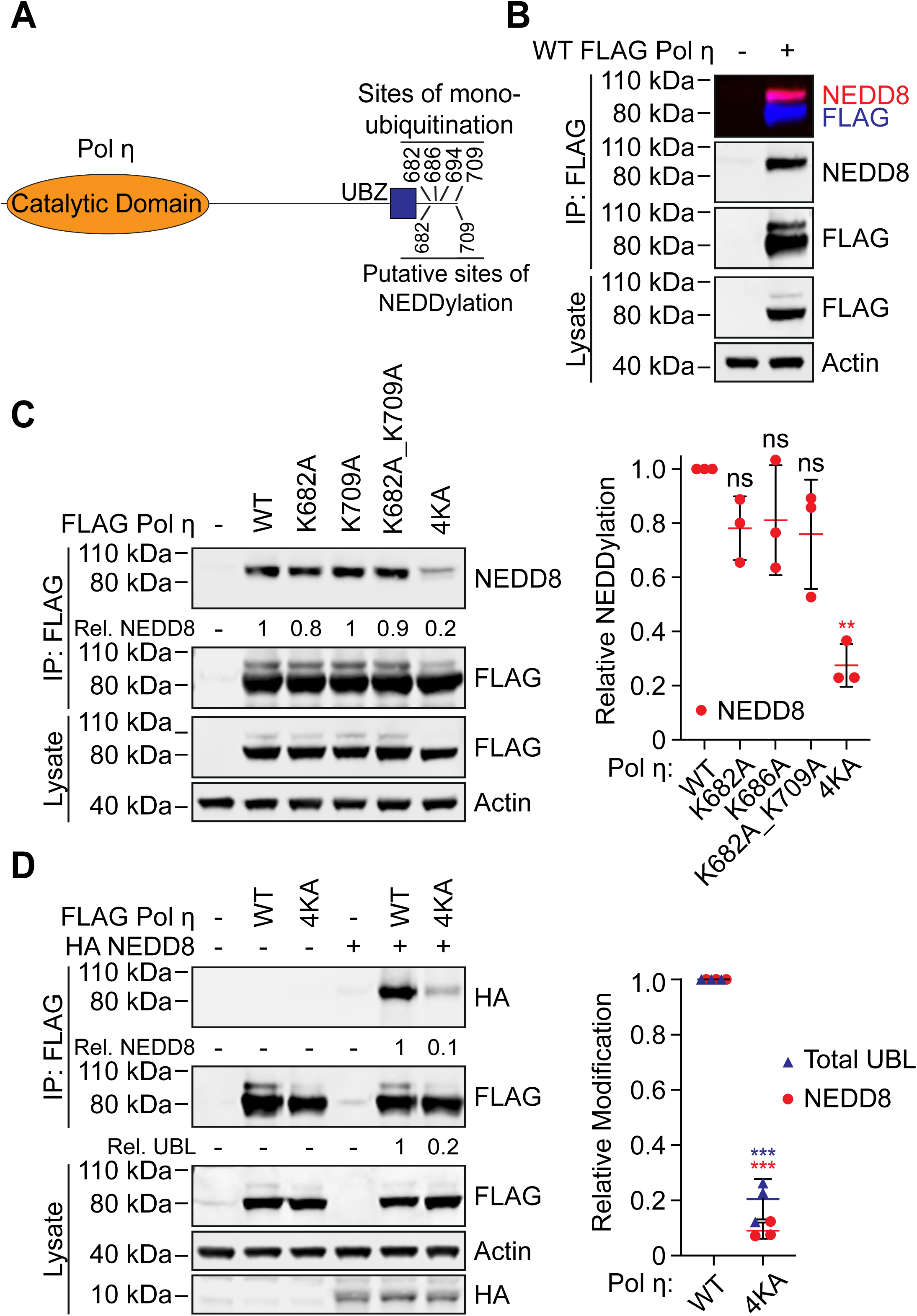
Pol η is a substrate of NEDD8. (**A**) Schematic of Pol η illustrating the location of lysine residues that can be mono-ubiquitinated, and putatively, mono-NEDDylated. (**B**) WT FLAG Pol η was immuoprecipitated from 293T cells and the lysate and eluent immunoblotted as indicated. FLAG and NEDD8 antibodies were detected with fluorescent secondary antibodies and visualized on separate channels. The top box shows a merged image of both channels. (**C**) WT and mutant FLAG Pol η was immunoprecipitated from 293T and the lysate and eluent immunoblotted as indicated. Relative NEDDylation was calculated based on the ratio of NEDD8 to FLAG in the eluent. The graph represents the quantification of relative NEDDylation from three independent repeats. Error bars represent standard deviation. Unpaired t-tests were used to assess differences in NEDDylation of WT vs mutant Pol η. ** = p < 0.1. (**D**) WT and 4KA FLAG Pol η was immunoprecipitated from 293T cells co-expressing HA NEDD8. The lysate and eluent were immunoblotted as indicated. Relative NEDDylation was calculated based on the ratio of HA to FLAG in the eluent. Relative UBLylation was calculated based on the ratio of higher to lower FLAG Pol η bands. The graph represents quantification from three independent repeats. Statistical analysis was performed as per (C). *** = p < 0.01.

To confirm that Pol η is a substrate of NEDD8, we immunoprecipitated FLAG-tagged Pol η from 293T cells, and immunoblotted eluting proteins with an antibody against NEDD8. This revealed a single major band that was cross-reactive with FLAG antibodies and migrated behind the predominant WT Pol η band (**Figure 1B**). Based on this migration pattern, we considered that this most likely represents the addition of a single NEDD8 molecule on Pol η. We also performed this experiment using forms of Pol η where we mutated K682 and K709, as well as two other C-terminal residues known to be mono-ubiquitinated (K686 and K694), to alanine. While the individual mutation of these residues, or the combined mutation of K682 and K709, had little effect on Pol η mono-NEDDylation, simultaneous mutation of all four lysine residues (4KA) reduced Pol η NEDDylation by ∼75 % compared to WT (**Figure 1C**). This is a similar level of reduction as has been reported for Pol η mono-ubiquitination following the combined mutation of these residues (16). Notably, while our data suggests that any of these lysine residues may be modified, the detection of only a single predominant mono-UBL band indicates that Pol η is only modified by a single UBL molecule at one time.

Ubiquitin and NEDD8 are structurally similar proteins sharing ∼60% primary sequence identity (32). We were therefore initially concerned whether the NEDD8 antibody could be non-specifically detecting ubiquitin in our samples. Indeed, despite the NEDD8 antibody having a strong preference for NEDD8, we did see a small degree of non-specific binding to high concentrations of recombinant ubiquitin in a dot blot assay (**Figure S1**). To detect Pol η NEDDylation by another approach, we therefore co-transfected 293T cells with FLAG Pol η and HA NEDD8. We then immunoprecipitated FLAG-tagged Pol η, and immunoblotted the eluted proteins with a HA antibody. This allowed us to readily and specifically detect mono-NEDDylation of WT Pol η, which was strongly reduced for 4KA Pol η (**Figure 1D**).

We nevertheless wanted to be assured that our NEDD8 antibody specifically detects Pol η mono-NEDDylation in our IP assays. To test this, we treated cells with the NEDDylation inhibitor, MLN4924 (33). This compound inhibits the NEDD8 activating enzyme (NAE), disrupting NEDD8 conjugation without affecting other UBL modifications. MLN4924 treatment reduced Pol η mono-NEDDylation by >95%, confirming the specificity of our detection method (**Figure 2A**). Interestingly, despite strongly reducing Pol η mono-NEDDylation, treatment with MLN4924 had little effect on total Pol η UBL levels. We interpreted this as mono-NEDDylation representing a minor portion of total steady-state Pol η modification by ubiquitin-like proteins (referred to hereafter as UBLylation).

**Figure 2:**
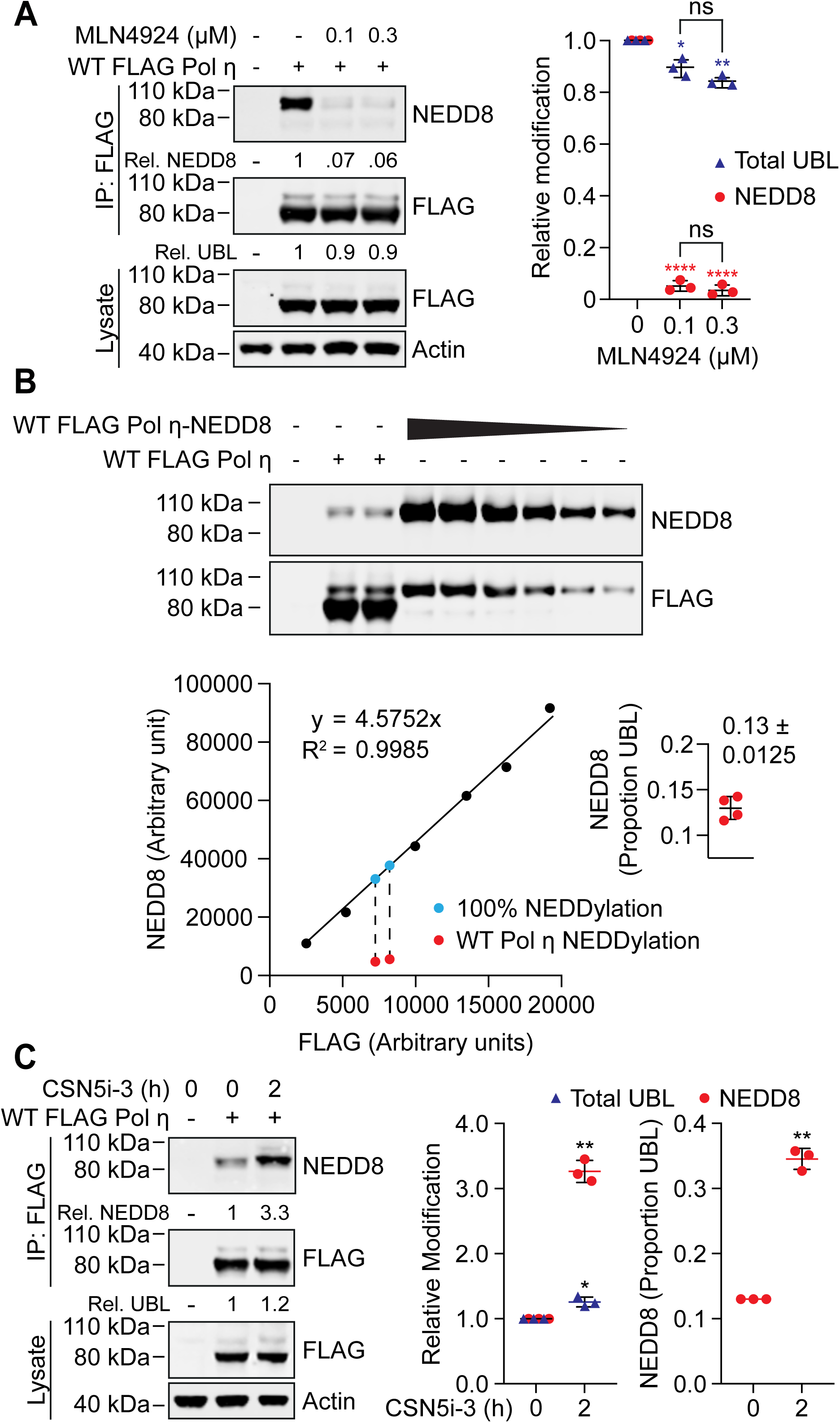
Pol η mono-NEDDylation is dynamically regulated in cells. (**A**) 293T cells expressing WT FLAG Pol η were treated with 0.1 or 0.3 μM MLN4924, or mock treated with DMSO, for 16 hours. WT FLAG Pol η was immunoprecipitated from these cells, and samples immunoblotted as indicated. Relative mono-UBLylation was calculated based on the ratio of higher to lower FLAG Pol η bands. Mono-NEDDylation was calculated based on the ratio of NEDD8 to FLAG bands. The graph represents calculated values from three independent experiments. Error bars represent standard deviation. Unpaired t-tests were used to assess differences in Pol η modification. * = p < 0.5, ** = p < 0.1, **** = p <0.001, ns = not significant. (**B**) WT FLAG Pol η, or a WT FLAG Pol η-NEDD8 chimera, was immunoprecipitated from 293T cells. A dilution series of the chimeric protein, and technical replicates of WT FLAG Pol η, were immunoblotted and probed with FLAG and NEDD8 antibodies. The FLAG and NEDD8 bands of the chimera were quantified and are represented by the graph. A standard curve defined by a linear equation was determined for each repeat, where the multiplication factor represents the average ratio of FLAG to NEDD8 for each dilution. The NEDD8 and upper FLAG bands of the two WT FLAG Pol η lanes were also quantified, as represented in red. The ratio of NEDD8 to FLAG for the WT protein was compared to that of the chimeric protein, to calculate NEDDylation as a proportion of total UBLylation. The calculated values from four technical repeats are depicted in the inset graph. (**C**) 293T cells expressing WT FLAG Pol η were treated with 1 μM of the COP9 inhibitor CSN5i-3, or DMSO for 2 hours. FLAG Pol η was immunoprecpitated from these cells and the eluent and lysate immunoblotted as indicated. Relative mono-NEDDylation and relative mono-UBLylation was determined as per (A) and is represented in the left graph. The right graph represents changes in NEDDylation as a proportion of total UBLylation, assuming steady state levels of 0.13, as calculated in (B). Statistical analysis was performed as per (A).

To determine this proportion more directly, we generated a Pol η-NEDD8 chimeric protein, where we fused NEDD8 to the C-terminus of FLAG Pol η. We then expressed and immunoprecipitated this chimera from 293T cells and immunoblotted a dilution series on a western blot next to immunoprecipitated WT Pol η (**Figure 2B**). This fusion protein allowed us to calculate a ratio of NEDD8 to FLAG signal that corresponds with 100% mono-NEDDylation and compared this to the NEDD8/FLAG ratio of UBLylated Pol η. Doing so allowed us to deduce that mono-NEDDylation constitutes ∼13% of Pol η mono-UBLylation in undamaged cells. We also noted that the Pol η-NEDD8 chimera was not further UBLylated, suggesting that even the in-line expression of NEDD8 at the Pol η C-terminus is sufficient to prevent further modification.

We next considered the possibility that Pol η may be NEDDylated to a higher degree than steady-state levels reveal, but that this modification might be actively suppressed by deNEDDylation. The COP9 signalosome is the predominant deNEDDylating enzyme complex in cells and is known to function in the DNA damage response pathway of nucleotide excision repair (34). We therefore treated cells with an inhibitor of CSN5 (35) – the catalytic subunit of COP9 – and probed for Pol η NEDDylation. This treatment resulted in a three-fold increase in Pol η NEDDylation following 2 hours of treatment (**Figure 2C**). Interestingly, while we detected an ∼20% increase in total mono-UBLylation, this is insufficient to account for the total increase in Pol η NEDDylation. i.e. given that NEDDylation comprises ∼13% of total mono-UBLylation, a three-fold increase in NEDDylation would be expected to increase total UBLylation by >40%. This suggests that while some of the increase in Pol η mono-NEDDylation is due to new mono-UBylation, the remainder likely results from a concurrent decrease in other UBL modifications.

### Mono-ubiquitination and mono-NEDDylation require a functional UBZ domain

Mono-ubiquitination of Pol η is dependent on ubiquitin-binding by its C-terminal ubiquitin-binding zinc finger (UBZ) domain; as such, mutations in the UBZ domain prevent Pol η mono-ubiquitination (10). Similar results have been observed for other ubiquitin-binding proteins, where the ubiquitin-binding domain is thought to interact with E2-laden ubiquitin molecules and promote E3-independent ubiquitination of nearby lysine residues (36). The covalently attached ubiquitin molecule is then positioned to interact intramolecularly with the UBZ domain, preventing binding to other free ubiquitin molecules (16,37). This model may explain how the Pol η UBZ domain promotes mono-ubiquitination of adjacent lysine residues (**Figure 3A**), as well as why Pol η is only modified by a single mono-UBL molecule at one time. As Pol η mono-NEDDylation occurs on the same residues as ubiquitination, and these modifications are mutually exclusive, we wondered whether NEDDylation might also be regulated by interactions between NEDD8 and the Pol η UBZ domain. Indeed, many of the key residues of ubiquitin that bind the UBZ domain are also present in NEDD8 (21) (**Figure S2**).

**Figure 3:**
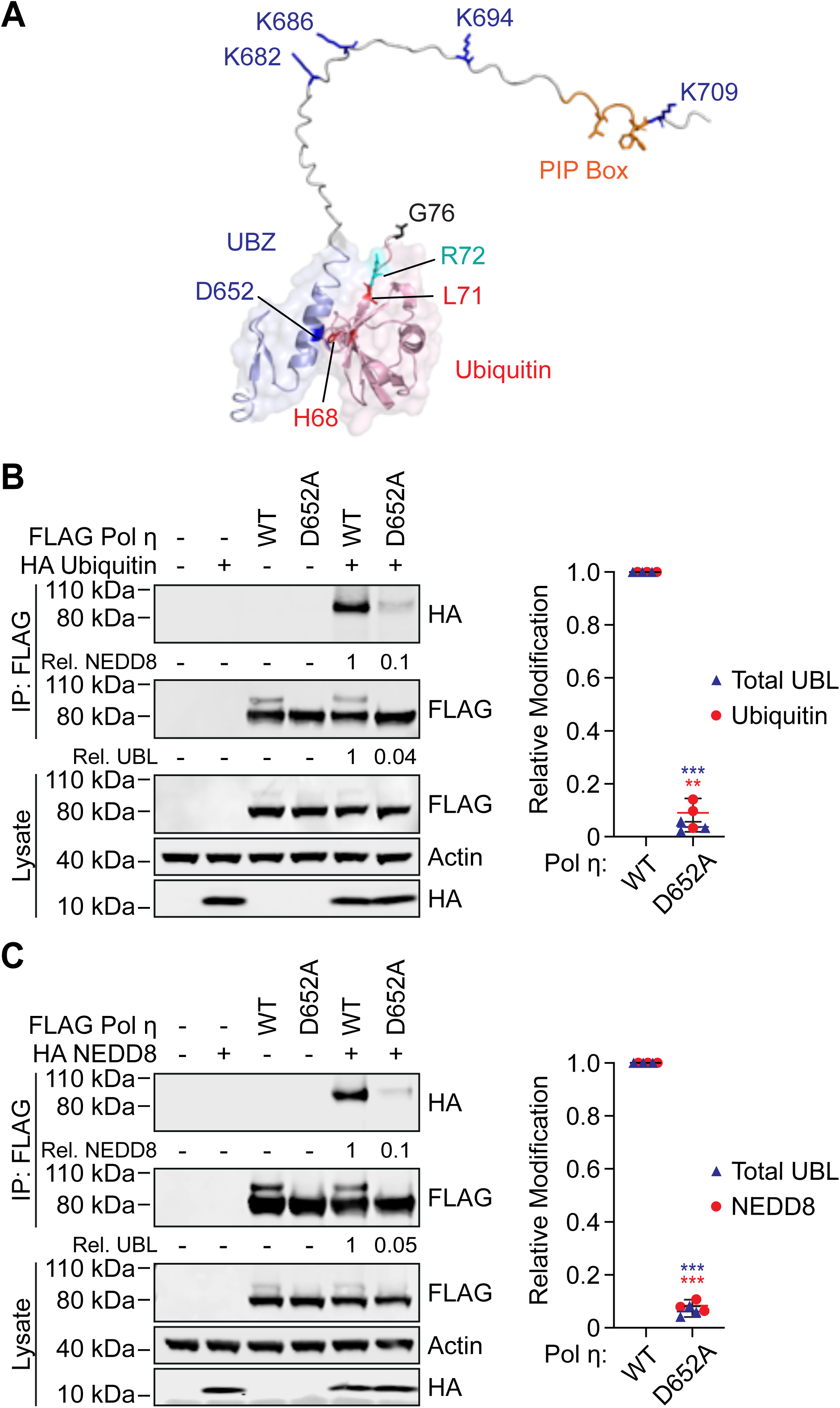
The UBZ domain of Pol η is required for mono-ubiquitination and mono-NEDDylation. (**A**) An AlphaFold generated model of ubiquitin in complex with the c-terminus of Pol η. (**B and C**) WT or D652A FLAG Pol η was immunoprecipitated from 293T cells co-expressing HA ubiquitin (B) or HA NEDD8 (C). The eluent and lysate were immunoblotted as indicated. Relative mono-UBLylation was calculated based on the ratio of higher to lower FLAG Pol η bands. Mono-ubiquitination and mono-NEDDylation was calculated based on the ratio of HA to FLAG bands. The graph represents calculated values from three independent experiments. Error bars represent standard deviation. Unpaired t-tests were used to assess differences in Pol η modification. * = p < 0.5, ** = p < 0.1, *** = p <0.01.

UBZ domains are specialized C_2_H_2_ zinc fingers consisting of two short β-strands and a C-terminal α-helix that forms a binding interface with the canonical hydrophobic surface of ubiquitin (10,11,21). To test our hypothesis, we introduced a previously described D652A mutation in the C-terminal α-helix of the Pol η UBZ domain (10) to prevent UBL binding and expressed this mutant in cells. D652A mutation disrupted mono-ubiquitination and mono-NEDDylation by ∼90% (**Figure 3B-C**). These data thereby support the notion that the UBZ domain is important for both forms of UBLylation.

We next sought to determine directly whether the UBZ domain binds to NEDD8. To do so, we purified isotopically labelled Pol η UBZ domain and used solution nuclear magnetic resonance (NMR) to monitor its binding to either unlabelled NEDD8 or ubiquitin. We then quantified the resulting chemical shift perturbations (CSPs) and mapped these onto the crystal structure of the UBZ domain (PDB: 3WUP) (**Figure 4A-B**). This revealed that NEDD8 and ubiquitin both bind to the same C-terminal region of the UBZ domain. While ubiquitin binding was notably tighter (K_D_=158.6 ± 9.5μM) than binding to NEDD8 (K_D_=671.3 ± 153.3μM) and involved larger chemical shift perturbations of UBZ residues e.g. F655 (**Figure 4C-D**), both interactions shared a binding interface that includes residue D652. This explains why D652A mutation disrupts both ubiquitination and NEDDylation of Pol η.

**Figure 4:**
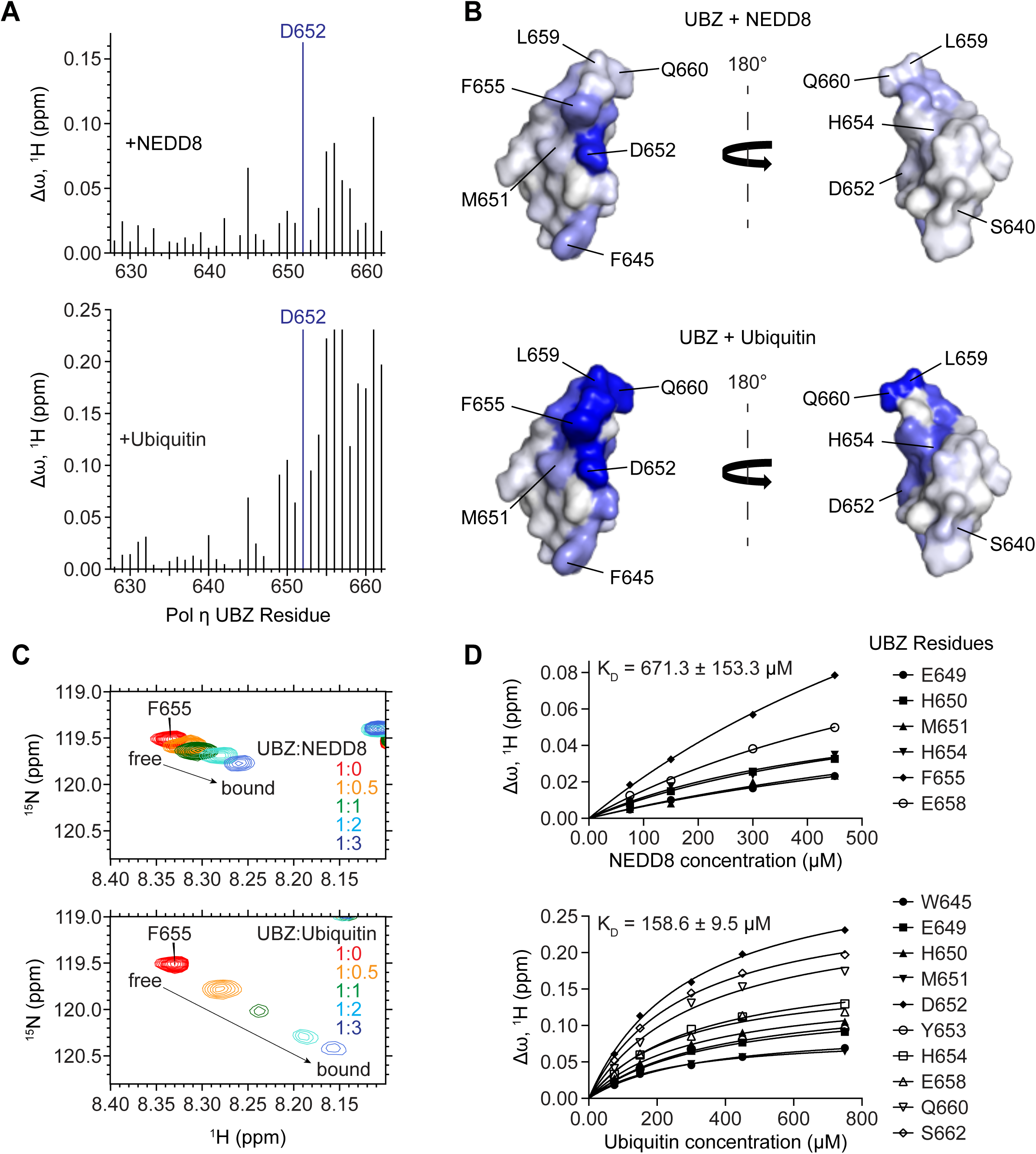
The UBZ domain of Pol η interacts with both ubiquitin and NEDD8 through a shared binding surface. (**A**) A bar graph of per-residue chemical shift perturbations (CSPs) in ^15^N-TROSY-HSQC spectrum of ^15^N-labeled Pol η UBZ domain caused by the addition of unlabelled NEDD8 (top panel) and ubiquitin (bottom panel). Residues for which peaks disappeared due to broadening were assigned to the highest observed CSP value. (**B**) NEDD8 (top) and ubiquitin (bottom) binding sites mapped on the surface of the UBZ (PDB: 3WUP). The surface is color-coded according to the NMR CSPs (Δω) from smallest (white) to largest (blue). The structures are shown in two orientations with a 180° rotation. (**C**) A representative region of the Pol η UBZ 15N-TROSY-HSQC spectra highlights the gradual transition of the peak corresponding to residue F655 from its free (red) to its bound conformation (blue) upon titration with either NEDD8 (top) or ubiquitin (bottom). (**D**) NMR titration plots used to estimate the K_D_ of binding.

We also used NMR to identify the UBZ-binding interface of NEDD8. In a reciprocal approach to the above, we incubated isotopically labelled NEDD8 with increasing concentrations of unlabelled UBZ domain, calculated chemical shift perturbations, and mapped these onto the NEDD8 crystal structure (PDB: 1NDD) to reveal a continuous binding interface (**Figure 5A-B**). NMR-derived interfacial residues of NEDD8 and UBZ were used to generate a structural model of the UBZ/NEDD8 complex using biomolecular docking (38,39) (**Figure 5C**). The resulting structure of the complex conclusively demonstrates the binding of the Pol η UBZ domain to the canonical hydrophobic surface of NEDD8, akin to the reported UBZ-ubiquitin interaction (27,40–42).

**Figure 5:**
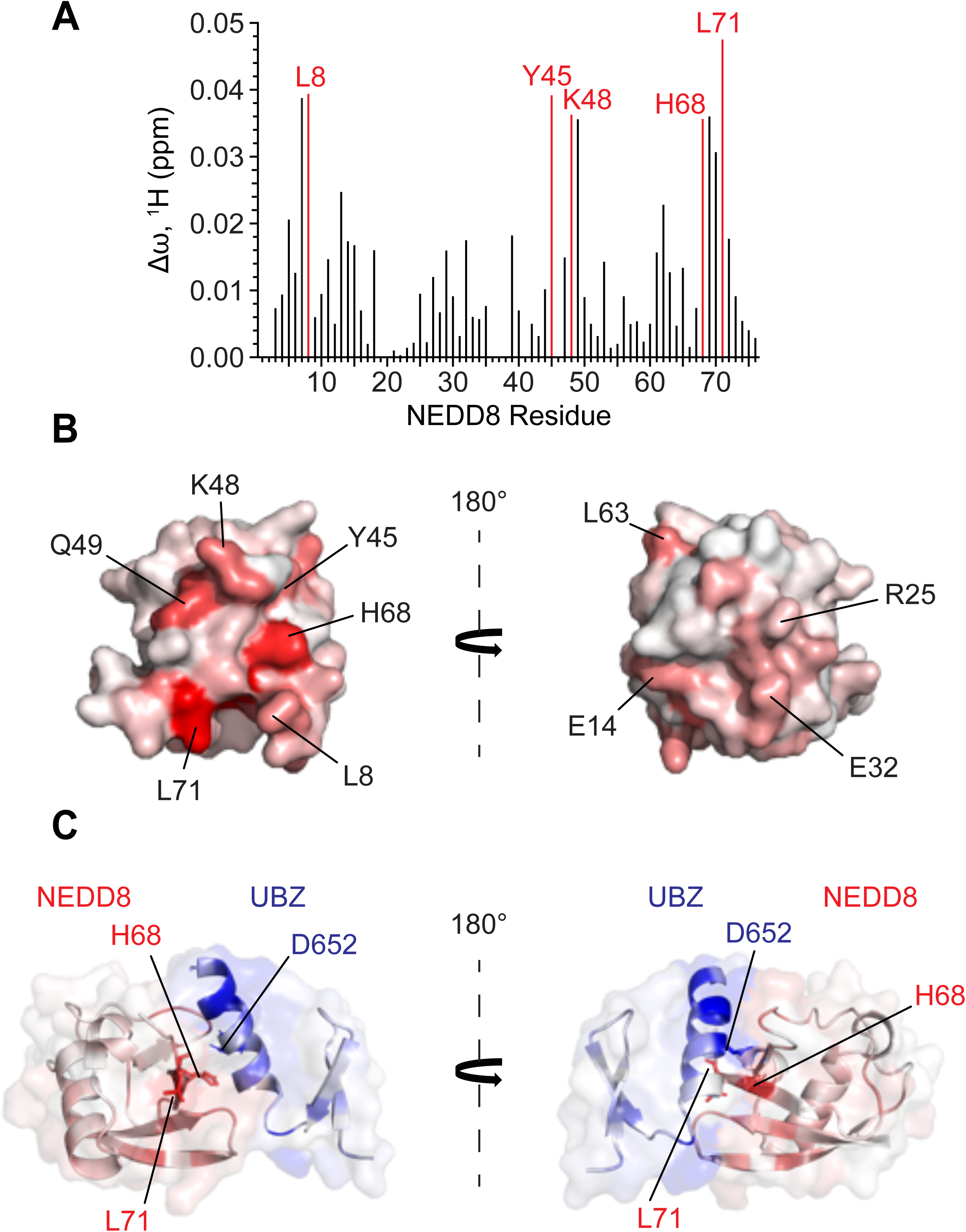
An NMR-based molecular model of the UBZ/NEDD8 complex. (**A**) A bar graph of per-residue chemical shift perturbations (CSPs) in ^15^N-TROSY-HSQC spectrum of ^15^N-labelled NEDD8 caused by the addition of unlabelled UBZ domain of Pol η. Residues for which peaks disappeared due to broadening were assigned to the highest observed CSP value. (**B**) The UBZ domain binding site mapped on the surface of NEDD8 (PDB: 1NDD). The surface is color-coded according to the NMR CSPs (Δω) from smallest (white) to largest (red). The structures are shown in two orientations with a 180° rotation (**C**) HADDOCK model of the UBZ domain of Pol η (blue) in complex with NEDD8 (red) colored according to their Δω as in Figure 4B and 5C. The model is shown in two orientations with a 180° rotation.

### Mono-ubiquitination and mono-NEDDylation differentially regulate Pol η localization to DNA damage sites

Our data above demonstrate that the C-terminus of Pol η can be modified by mutually exclusive mono-ubiquitination or mono-NEDDylation. It was however unclear how these modifications might regulate Pol η function. Mono-ubiquitination has a well-established role in regulating the assembly of TLS complexes at sites of damage. We therefore wondered whether ubiquitination and NEDDylation might differentially regulate DNA damage-induced Pol η localization.

To test this, we expressed eGFP-tagged Pol η in MRC-5 cells and monitored UVC-induced DNA damage foci formation by fluorescent imaging. In addition to WT and 4KA Pol η, we expressed variants containing mutations in the UBZ domain (D652A), or the C-terminal PIP box (L704A_F707A_F708A), to disrupt ubiquitin- and PCNA-binding, respectively. We also created chimeric proteins where we fused either ubiquitin or NEDD8 to the C-terminus of Pol η, which we used to mimic constitutive mono-ubiquitination and mono-NEDDylation, respectively. WT Pol η readily formed DNA damage-induced foci, as did the Pol η-ubiquitin chimera. Interestingly, the 4KA mutant and NEDD8-fusion protein both failed to do so, resembling variants containing mutations in the PIP box or UBZ domain (D652A) (**Figure 6A**). We considered that one explanation for these data might be that mono-ubiquitination is important for the accumulation of Pol η at foci. This may also explain the lack of foci formation for the NEDD8-fusion protein, as similar to native lysine mono-NEDDylation, we observed that fusion to NEDD8 prevents further Pol η mono-ubiquitination (Figure 2B). This model is however inconsistent with previous work, which suggested that Pol η mono-ubiquitination could inhibit binding to other ubiquitinated proteins (16). To test whether UBLylated Pol η can associate with mono-ubiquitinated PCNA, we used immunoprecipitation assays to examine Pol η binding to WT PCNA, K164R PCNA, or a chimera where we fused ubiquitin to the C-terminus of K164R PCNA. A similar PCNA-ubiquitin chimera has been described previously and been found to effectively mimic mono-ubiquitinated PCNA and support DNA damage tolerance in *S. cerevisiae* (43). The effectiveness of this mimic is likely due to the proximity of PCNA K164 to its C-terminus, as well as the flexibility of the PCNA C-terminal residues, and the C-terminus of Pol η (**Figure S3**). In these assays, mono-UBLylated and non-UBLylated Pol η immunoprecipitated with the PCNA-ubiquitin fusion protein (**Figure 6B**). Although we cannot distinguish mono-ubiquitinated from mono-NEDDylated Pol η in these assays, these data indicate that mono-UBLylation does not exclude Pol η from binding to other ubiquitinated proteins.

**Figure 6:**
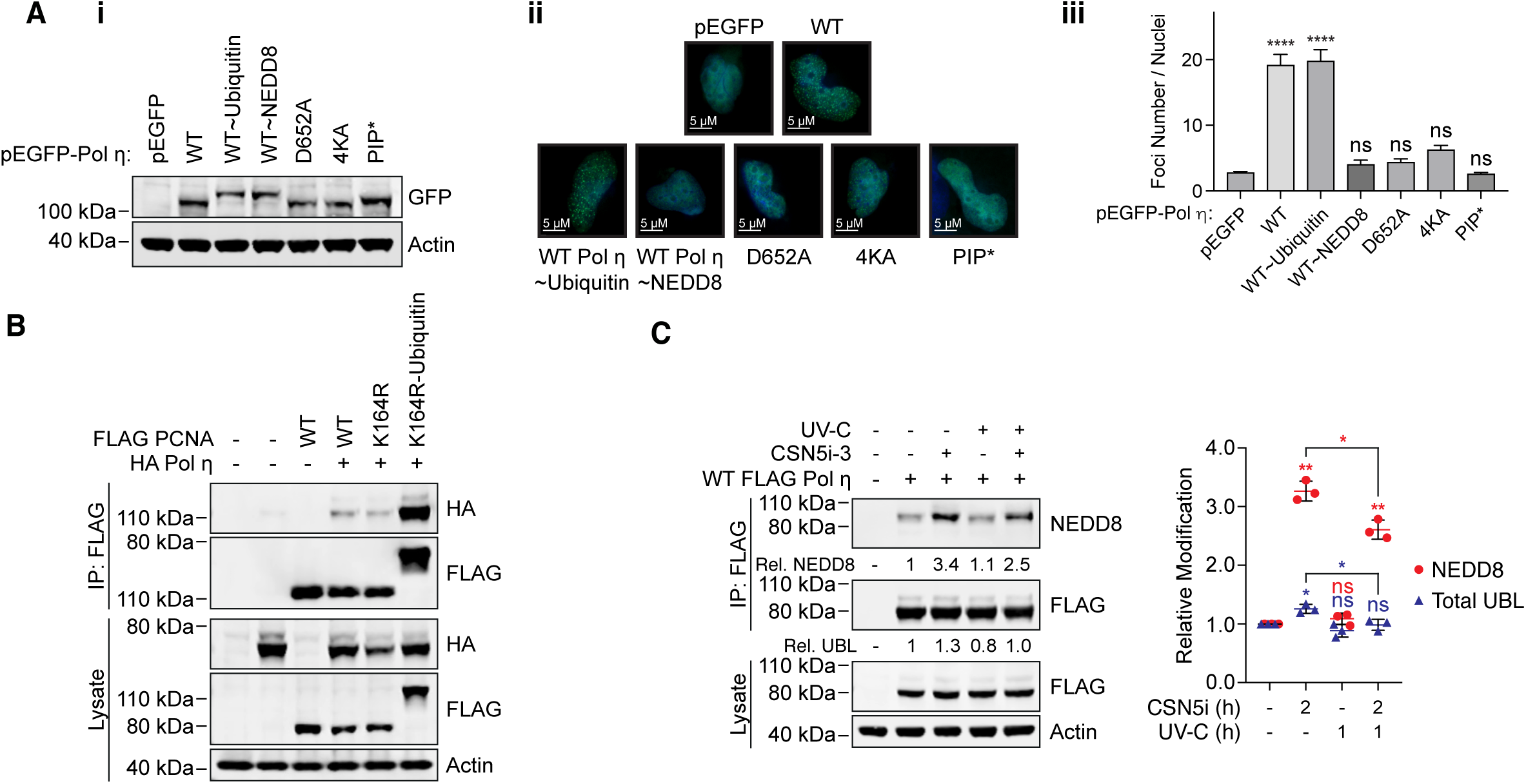
Mono-ubiquitination and mono-NEDDylation differentially regulate Pol η. (**A**) The indicated GFP-Pol η variants were transiently expressed in MRC-5 cells. (i) The western blot demonstrates expression of each variant in whole cell lysates. (ii) Cells expressing each construct and growing on coverslips were exposed to 20 Jm² of UV-C, fixed after 6 hours, and imaged on a fluorescence microscope with a 63x objective lens. Images taken in the EGFP channel were converted to 16-bit grayscale and the number of foci per nucleus measured. (iii) The graph represents the average number of foci per nucleus. Error bars represent standard deviation. Unpaired t-tests were used to assess differences between groups. **** = p <0.001, ns = not significant. (**B**) WT, K164R and a K164R-ubiquitin chimera of PCNA were immunoprecipitated from cells co-expressing HA Pol η. The lysate and eluent were immunoblotted as indicated. (**C**) 293T cells expressing WT FLAG Pol η were treated with 1 μM of the NEDDylation inhibitor CSN5i-3, or DMSO for 1 hour, before being exposed to 20 Jm² of UV-C. Cells were harvested 1 hour after exposure, FLAG Pol η was immunoprecipitated, and the eluent and lysate immunoblotted as indicated. Relative mono-UBLylation was calculated based on the ratio of higher to lower FLAG Pol η bands. Mono-NEDDylation was calculated based on the ratio of NEDD8 to FLAG bands. The graph represents quantification from three independent repeats. Error bars represent standard deviation. Unpaired t-tests were used to assess differences in NEDDylation and UBLylation. * = p < 0.05, ** = p < 0.1, ns = not significant.

We also reasoned that if NEDDylation disrupts the accumulation of Pol η at DNA damage sites, levels of this modification may be altered in response to DNA damage. To test this, we exposed cells to UV-C in the presence or absence of the COP9 inhibitor, immunoprecipitated Pol η and immunoblotted for NEDD8. While UV-C exposure had minimal effects on steady state NEDDylation levels, we observed a small but significant reduction in NEDDylation when cells were co-treated with the COP9 inhibitor (**Figure 6C**). Together, these results support that mono-NEDDylation negatively regulates Pol η, likely by disrupting mono-ubiquitination.

Taken together, our results demonstrate that Pol η is a substrate of NEDD8 and its NEDDylation is directed to the C-terminus by the UBZ domain. Furthermore, the NEDDylation of Pol η is mutually exclusive with its ubiquitination and negatively regulates Pol η accumulation at sites of DNA damage.

## DISCUSSION

Pol η is a ubiquitin-binding protein that mediates its own mono-ubiquitination at four C-terminal lysine residues – K682, K686, K694 and K709 (10,16,17,21). In this work, we demonstrate that the UBZ domain of Pol η can also bind NEDD8 to facilitate mono-NEDDylation of the same sites. While other types of ubiquitin-binding domains have been shown to interact with NEDD8 (44), this is the first example involving a UBZ-type domain. This interaction occurs in a manner analogous to ubiquitin-binding, involving the helix of the UBZ domain, and the canonical UBL hydrophobic surface of NEDD8. Discriminatory binding to ubiquitin over NEDD8 is often dictated via a requirement of ubiquitin-binding domains to interact electrostatically with ubiquitin residue R72 (A72 in NEDD8) (28,45,46). An NMR-based study of ubiquitin-binding by the Pol η UBZ domain however found that ubiquitin R72 was not significantly perturbed upon complex formation, suggesting this residue is not part of the Pol η UBZ-binding interface (21) (**Figure 3 and S2**). This may provide some explanation for why Pol η does not strictly discriminate between interacting with ubiquitin and NEDD8. Although we did find the UBZ domain bound ubiquitin more tightly, this is likely due to other variations in the hydrophobic surfaces of these UBLs.

Ubiquitin-binding is thought to mediate Pol η mono-ubiquitination by recruiting ubiquitin-laden E2 enzymes, as has been described for other mono-ubiquitinated ubiquitin-binding proteins (36). As such, mutations in the UBZ domain disrupt mono-ubiquitination (16). Here we find that mutating the central UBZ residue, aspartate 652, to alanine, also prevents mono-NEDDylation, suggesting these modifications are regulated in an analogous manner. Although NEDD8 and ubiquitin are found at similar concentrations in cells (47), weaker binding to NEDD8 could explain why mono-NEDDylation accounts for a smaller portion of total UBLylation at steady-state levels, potentially due to lower rates of recruitment of NEDD8-E2 complexes. Our observations also support that Pol η mono-ubiquitination or mono-NEDDylation prevents further UBLylation at other lysine residues. This is apparent from the presence of a single predominant higher molecular weight Pol η species on immunoblots. Previous works have proposed that when Pol η is mono-ubiquitinated, its UBZ domain may interact intramolecularly with the ubiquitin molecule, preventing the UBZ domain from mediating the mono-ubiquitination of adjacent lysine residues (16,17). It is possible that Pol η mono-NEDDylation prevents further UBLylation due to similar intramolecular interactions between the UBZ domain and NEDD8. Indeed, weaker binding of the UBZ with NEDD8 versus ubiquitin may permit greater access of deNEDDylating enzymes, allowing for the strong negative regulation of mono-NEDDylation evident in cells treated with a COP9 inhibitor. Interestingly, we did not observe additional mono-UBLylation of a Pol η-NEDD8 chimera, suggesting that even the non-native fusion of NEDD8 to the Pol η C-terminus – rather than linkage of NEDD8 G76 to a Pol η lysine residue – is sufficient to prevent further modification. This may be due to the flexibility of the Pol η C-terminus, allowing the NEDD8 molecule to interact with the UBZ domain in a range of confirmations.

In addition to recruiting ubiquitin-E2 complexes, the Pol η UBZ domain also permits binding to other ubiquitinated proteins, most notably PCNA mono-ubiquitinated at K164. Interestingly, this residue of PCNA can also be mono-NEDDylated, which has been suggested to disrupt translesion synthesis by antagonizing mono-ubiquitination and the accumulation of TLS polymerases (48). Although we conclusively demonstrate that Pol η can bind NEDD8, it is conceivable that the weaker binding we observe is insufficient to allow Pol η recruitment in this context.

Previous works have proposed that intramolecular UBZ-ubiquitin interactions may not only limit mono-ubiquitinated Pol η from interacting with ubiquitin-E2 complexes, but also prevent Pol η from interacting with other ubiquitinated proteins (16,17). Our immunoprecipitation data, however, demonstrated that mono-UBLylated Pol η can associate with a PCNA-ubiquitin chimera. Furthermore, we found that a Pol η-ubiquitin fusion protein readily localizes to DNA damage-induced foci (**Figures 6A and B**). This suggests that the intramolecular UBZ-ubiquitin interactions may be disrupted in some contexts. In the case of binding to mono-ubiquitinated PCNA, this is likely due to the additional stabilization afforded by the Pol η PIP box-PCNA interaction and positioning of the UBZ domain near the ubiquitin group on PCNA (49). It is also possible that the interaction between mono-ubiquitinated Pol η and mono-ubiquitinated PCNA is stabilized by additional proteins that interact with the ubiquitin group on Pol η.

Our observation that 4KA mutation disrupts Pol η from forming DNA damage-induced foci is consistent with a previous study (16). In their work, however, the authors suggested that these UBLylated lysine residues reside within an extended PCNA-binding interface of Pol η, that includes not only the PIP box (M701 – F708), but also additional residues within the upstream nuclear localization signal (K682 – K694) (**Figure 7A**). The presence of an extended PCNA-binding interface within Pol η has not however been supported experimentally. Indeed, crystal structures of PCNA bound to a PIP box-containing Pol η peptide (K694 – H713), demonstrated that PCNA does not interact with Pol η K694, or other Pol η amino acids N-terminal to M701 (50). In addition, while K709 of Pol η formed backbone hydrogen bonds with PCNA, its side chain was extended away from PCNA (**Figure S3**). AlphaFold modelling of an interaction between PCNA and Pol η furthermore supports that amino acids L657-M701 of Pol η (between the UBZ domain and the PIP box) comprise an unstructured and flexible region, which is likely important for the Pol η PIP box to bind the surface of PCNA and permit the UBZ domain to simultaneously bind ubiquitinated K164 (**Figure 7B**). It is therefore unlikely that Pol η forms an extended binding interface with the surface of PCNA outside of the defined PIP-PCNA interacting region, or that lysine residues in this region of Pol η directly interact with PCNA.

**Figure 7:**
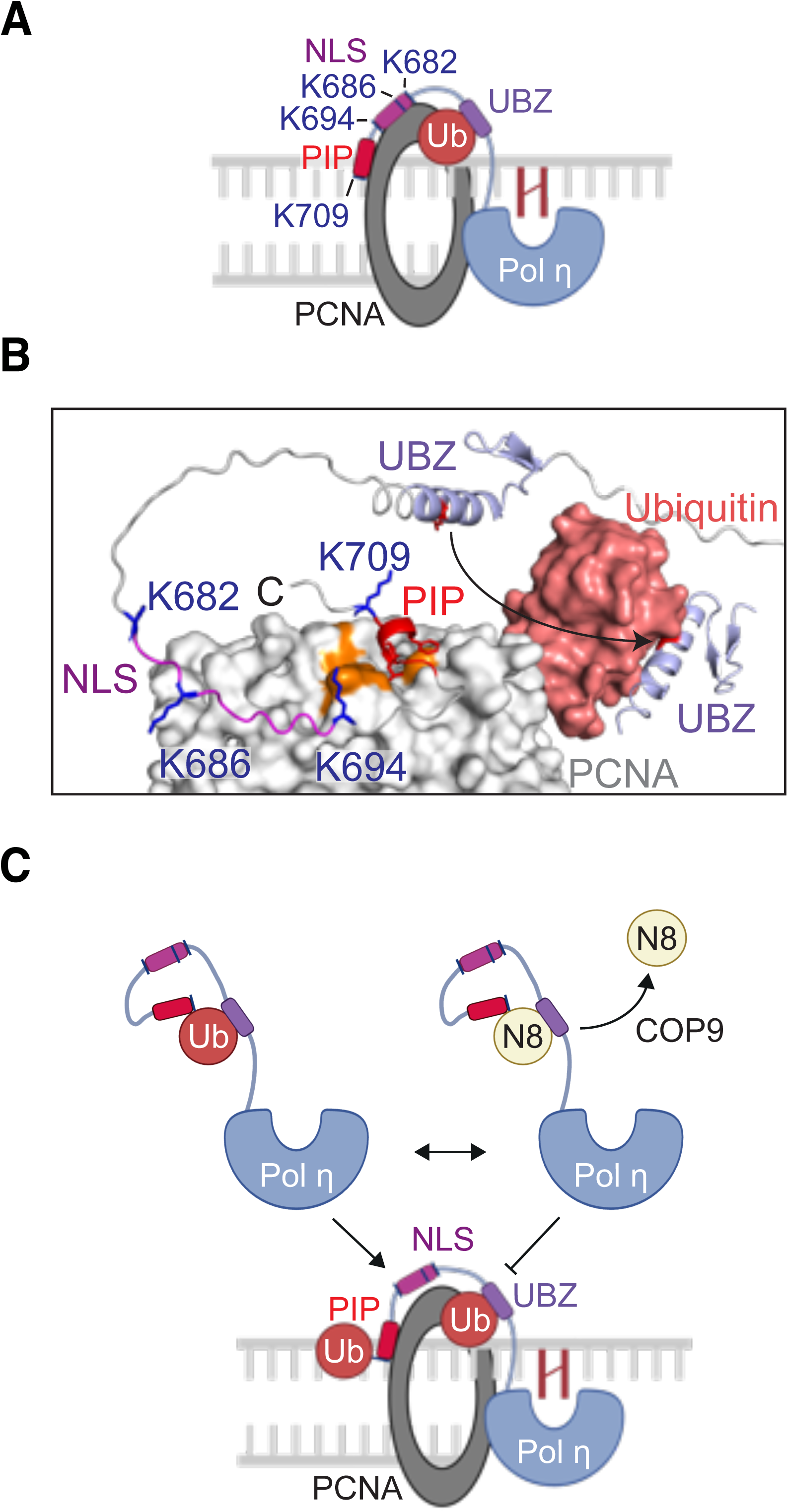
Models for the regulation of Pol η by mono-UBLylation. (**A**) A schematic illustrating a previously proposed model for the interaction between Pol η and mono-ubiquitinated PCNA (16). The PIP box and nuclear localization signal (NLS) of Pol η were suggested to form an extended binding interface with PCNA, while the ubiquitin-binding zinc finger (UBZ) binds the mono-ubiquitin moiety (**B**) An AlphaFold 3 model of the Pol η C-terminus in complex with PCNA. This model was overlaid with the crystal structure of mono-ubiquitinated PCNA (PDB 3TBL) to position the ubiquitin group. AlphaFold was also used to model the UBZ-PCNA interaction. The orange surface of PCNA represents the universal binding site, which can be seen interacting with the Pol η PIP box. (**C**) A model of Pol η regulation by mono-ubiquitination and mono-NEDDylation.

We therefore suggest an alternative explanation – that Pol η mono-ubiquitination is required for the proper accumulation of Pol η at sites of DNA damage (**Figure 7C**). This may explain why both 4KA Pol η and a Pol η-NEDD8 fusion proteins fail to form foci, as neither variant is readily ubiquitinated. Although not determined here, one explanation for this difference may be that mono-ubiquitination allows Pol η to interact with other ubiquitin-binding proteins that do not bind NEDD8. The identity of these interactors will however require further study. Nevertheless, we suggest that mono-NEDDylation of Pol η represents a negative regulatory mechanism that competitively disrupts the mono-ubiquitination of Pol η lysine residues. Our work thereby proposes a new mode for the regulation of Pol η by ubiquitin-like proteins

## DATA AVAILABILITY

Uncropped immunoblot images, as well as raw microscopy data, are available from Mendeley Data (https://data.mendeley.com/datasets/yrzngz38g3). All raw NMR data is available upon request. All expression vectors created by our labs have been deposited in the Addgene public plasmid repository (https://www.addgene.org/Roger_Woodgate/).

*Please note* the Mendeley data link is not yet active. A temporary pre-publication link is available here: https://data.mendeley.com/datasets/yrzngz38g3/draft?a=dd86dc41-2166-4d2c-9bec-a2b76cb3e68f

## SUPPLEMENTARY DATA

Supplementary Data are available at NAR online.

## AUTHOR CONTRIBUTIONS

**Natália Cestari Moreno:** Conceptualization, Methodology, Investigation, Formal analysis, Writing – Original Draft, Supervision. **Emilie J. Korchak:** Investigation, Formal analysis. **Marcela Teatin Latancia:** Investigation, Formal analysis. **Dana D’Orlando:** Investigation. **Temidayo Adegbenro:** Investigation. **Irina Bezsonova:** Investigation, Formal analysis, Writing – Review & Editing, Supervision. **Roger Woodgate:** Project Administration, Supervision, Writing – Review & Editing. **Nicholas Ashton**: Conceptualization, Investigation, Formal analysis, Writing – Original Draft, Writing – Review & Editing, Supervision.

## ACKNOWLEDGEMENTS

We acknowledge use of the web portals from the FP7 WeNMR, H2020 West-Life, EOSC-hub, and EGI-ACE European e-Infrastructure projects, and recognize their supporting organizations and national GRID Initiatives, for their vital contributions to the EGI infrastructure. We also thank Professor Pei Zhou (Duke University Medical Center) for the Pol η UBZ expression vector.

