## Supplemental Data for "DNA polymerase η is regulated by mutually exclusive mono-ubiquitination and mono-NEDDylation"

**Supplemental Data for: DNA Polymerase η is regulated by competitive mono-ubiquitination and mono-NEDDylation**

**This document includes:**

Supplemental Materials and Methods

Table S1

Figures S1 to S4

References for Supplemental Information

### Supplemental Materials and Methods

#### Dot blot

Recombinant human NEDD8 (R&D Systems #UL-812-500) or ubiquitin (R&D Systems #U-100H-10M) was diluted to 10 µg µL$^{-1}$ in sample buffer (50 mM Tris pH 8.0, 150 mM NaCl and 0.1 mM EDTA) to prepare a stock, then further diluted to concentrations of 2, 1, 0.5, and 0.25 µg µL$^{-1}$. 0.5 µL of sample was spotted onto a nitrocellulose membrane and allowed to air dry for 30 minutes. Proteins were visualized with a reversible total protein stain (Revert total protein stain, LiCor #926-11011) and imaged on an Odyssey CLX infrared imaging system (Li-Cor) at 700 nM. Membranes were then destained, blocked, and immunoblotted with ubiquitin (E4I2J; Cell Signaling Technology #43124) or NEDD8 antibodies (E19E3; Cell Signaling Technology #2754). Primary antibodies were detected using IRDye 800CW-conjugated anti-rabbit fluorescent secondary antibodies (Li-Cor) and visualized at 800 nM.

**Supplemental Table S1.** Expression constructs used in this study

| Plasmid | Mammalian/ E. coli | Figure(s) | Source | Addgene # |
|---|---|---|---|---|
| pCMV6-AN-DDK_ WT POLH (pJRM160) | Mammalian | 1B-D, 2A-C, 3B-C, 6C | (1) | 221897 |
| pCMV6-AN-DDK_ K682A POLH (pNCM23) | Mammalian | 1C | This work | 221862 |
| pCMV6-AN-DDK_ K709A POLH (pNCM24) | Mammalian | 1C | This work | 221863 |
| pCMV6-AN-DDK_ K682A_K709A POLH (pNCM25) | Mammalian | 1C | This work | 221864 |
| pCMV6-AN-DDK_ K682A_K686A_K694_K709A POLH (4KA) (pNCM26) | Mammalian | 1C | This work | 221865 |
| pcDNA3.1(+)-N-HA _HA NEDD8 (pNCM18) | Mammalian | 1D, 3C | This work | 221859 |
| pCMV6-AN-DDK_ WT POLH_ΔGG NEDD8 (pNCM21) | Mammalian | 2B | This work | 221860 |
| pCMV6-AN-DDK_ D652A POLH (pNWA8) | Mammalian | 3B, 3C | This work | 222006 |
| pCMV6-AN-HA_ Ubiquitin (pJRM147) | Mammalian | 3B | (2) | 131258 |
| pEGFP-C1_NLS (pNCM36) | Mammalian | 6A | This work | 221867 |
| pEGFP-C1_NLS_ WT POLH (pNCM37) | Mammalian | 6A | This work | 221868 |
| pEGFP-C1_NLS_ WT POLH_ΔGG Ubiquitin (pNCM38) | Mammalian | 6A | This work | 221869 |

| | | | | |
|---|---|---|---|---|
| pEGFP-C1_NLS_<br>WT POLH_∆GG NEDD8<br>(pNCM39) | Mammalian | 6A | This work | 221870 |
| pEGFP-C1_NLS_<br>D652A POLH<br>(pNCM40) | Mammalian | 6A | This work | 221871 |
| pEGFP-C1_NLS_<br>K682A_K686A_K694_K709A<br>POLH (4KA)<br>(pNCM41) | Mammalian | 6A | This work | 221872 |
| pEGFP-C1_NLS_<br>L704A_F707A_F708A<br>POLH (PIP)<br>(pNCM42) | Mammalian | 6A | This work | 221873 |
| pcDNA3.1(+)-N_DYK_<br>WT PCNA<br>(pNCM47) | Mammalian | 6B | This work | 221858 |
| pcDNA3.1(+)-N_DYK_<br>K164R PCNA<br>(pNCM48) | Mammalian | 6B | This work | 221874 |
| pcDNA3.1(+)-N_DYK_<br>K164R PCNA_∆GG Ubiquitin<br>(pNCM49) | Mammalian | 6B | This work | 221875 |
| pCMV6-AN-HA_<br>WT POLH<br>(pJRM56) | Mammalian | 6B | (3) | 201671 |
| pET15b_Pol η UBZ | E. coli | 4A-D<br>5A-B | (4) | - |
| pET-15b_Ubiquitin | E. coli | 4A-D | (5) | - |
| pET-28b(+)_N-His_NEDD8<br>(pNCM35) | E. coli | 4A-D<br>5A-B | This work | 221866 |

### Supplemental Figures

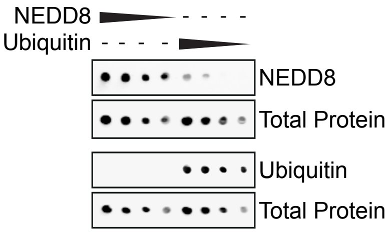


**Figure S1**: **Specificity of detection of the NEDD8 antibody**. A dot blot of recombinant human NEDD8 or ubiquitin (0.5 µL of 2, 1, 0.5, or 0.25 µg $\mu L^{-1}$ of protein). Membranes were stained to detect total protein, then immunoblotted with antibodies against NEDD8 or ubiquitin.

---

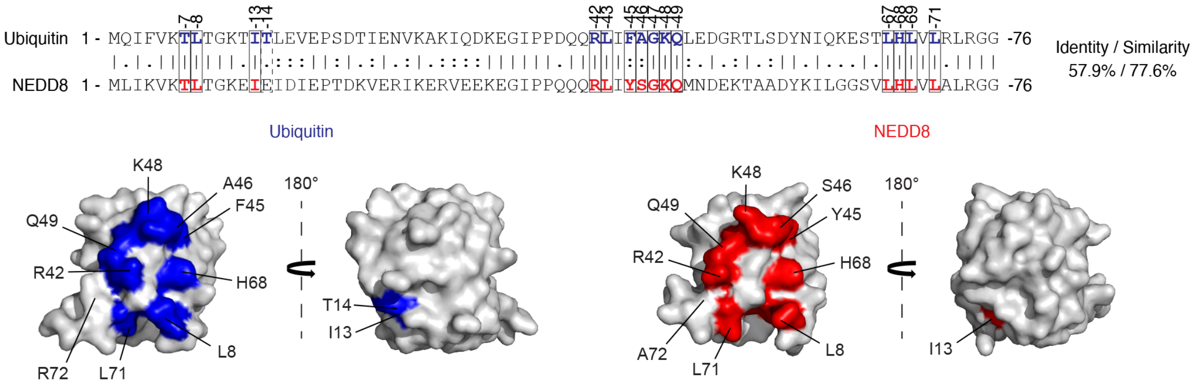


**Figure S2**: **The UBZ-binding residues of ubiquitin are conserved in NEDD8**. An alignment of the ubiquitin and NEDD8 primary sequences. The blue residues of ubiquitin are those which have previously been shown to be form the UBZ-binding surface (4). The corresponding residues of NEDD8 are shown in red where these residues are identical or similar. These UBZ-binding and corresponding residues are highlighted on the crystal structures of ubiquitin (PDB:1ubq) (6) or NEDD8 (PDB:1ndd) (7).

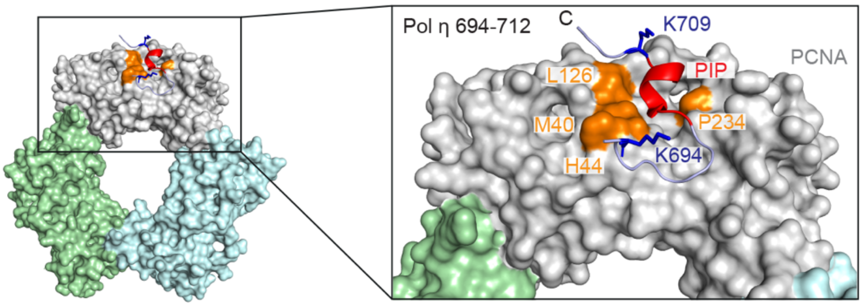


**Figure S3: The sidechains of Pol η K694 and K709 do not interact with PCNA**. A published crystal structure of a PIP-box containing Pol η peptide (amino acids 694-712) in complex with PCNA (PDB: 2zvk) (8) revealed that the side chains of K694 and K709 are directed away from the PCNA surface. The PCNA residues highlighted in orange define the PIP-binding universal binding site of PCNA.

---

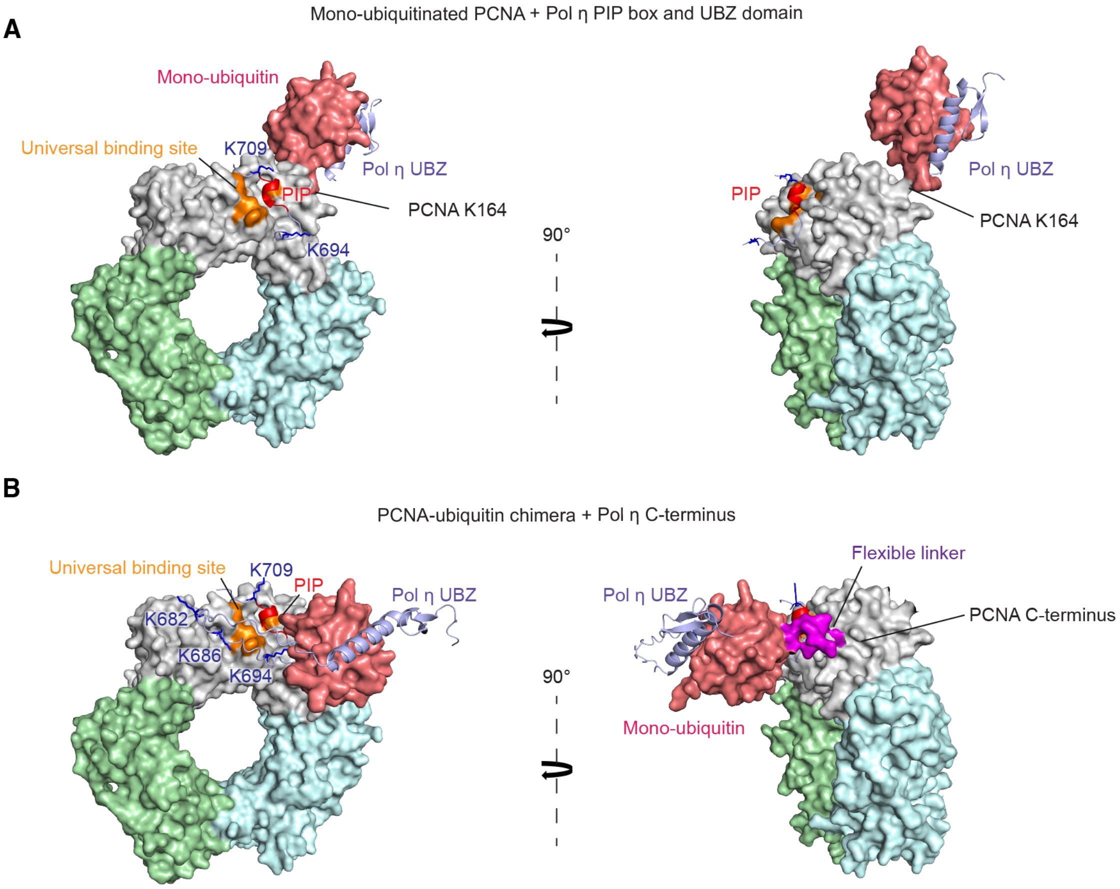

**Figure S4: A PCNA-ubiquitin chimera mimics mono-ubiquitinated PCNA** (**A**) A model of mono-ubiquitinated PCNA in complex with the PIP box (amino acids 694-712) and UBZ (amino acids 625-664) of Pol η. This model was assembled from a crystal structure of mono-ubiquitinated PCNA (PDB: 3tbl) (9), a crystal structure of the Pol η PIP box in complex with PCNA (PDB: 2zvk) (8), and an AlphaFold 3 (10) model of the Pol η UBZ domain bound to ubiquitin. (**B**) An AlphaFold model of the PCNA-ubiquitin chimera in complex with the C-terminus of Pol η (amino acids 634-713)
